# Intravital quantification of absolute cytoplasmic B cell calcium reveals dynamic signaling across B cell differentiation stages

**DOI:** 10.1101/2019.12.13.872820

**Authors:** Carolin Ulbricht, Ruth Leben, Asylkhan Rakhymzhan, Frank Kirchhoff, Lars Nitschke, Helena Radbruch, Raluca A Niesner, Anja E Hauser

## Abstract

Development, function and maintenance of lymphocytes largely depend upon the cellular mobilization and storage of Ca^2+^ ions. In B lymphocytes, the absolute amount of calcium mobilized and retained after cell signaling remains unknown, athough it is a crucial part of their selection within germinal centers and differentiation into plasma cells. Here, we introduce the novel reporter mouse strain YellowCaB that expresses the genetically encoded calcium indicator TN-XXL in CD19^+^ lymphocytes. The construct consists of the electron-donor fluorophore eCFP and the acceptor citrine, linked by a calcium sensitive domain. Its conformation and therefore donor quenching is directly linked to cytosolic calcium concentrations. By combining intravital two-photon fluorescence lifetime microscopy with our numerical approach for phasor-based analysis, we are able to extract absolute cytoplasmic calcium concentrations in activated B cells for the first time *in vivo*. We show that calcium concentrations in B cells are highly dynamic and fluctuations persist in extrafollicular B cells with functional relevance.

## Introduction

The capacity of the immune system to produce a variety of different antibodies and to further fine-tune their affinity to bind antigen (AG) upon pathogen challenge is one of the pillars of adaptive humoral immunity. Fine-tuning is achieved by somatic hypermutation of immunoglobulin genes, followed by a T cell-aided selection process, which B cells undergo within germinal centers (GC). This eventually results in a pool of high affinity memory B cells and long-lived plasma cells^1–3^. B cells encounter and take up membrane-bound AG in the form of immune complexes on follicular dendritic cells (FDCs) in GCs. This leads to B cell receptor (BCR) activation and calcium influx into the cell^4^. This eventually switches on effector proteins and transcription factors like nuclear factor kappa B (NF-κB), nuclear factor of activated T cells (NFAT), or Myelocytomatosis oncogene cellular homolog (c-Myc), thereby activating the B cells, inducing differentiation events and remodeling of metabolic requirements^5–10^. BCR-affinity dependent AG-capture has been thought to serve solemnly the processing and MHCII-dependent presentation to follicular helper T cells and that signaling is dampened^11^. However, newer studies show that BCR activated calcium signaling has to precede T cell derived signals and that the latter have to occur within a limited period of time after initial BCR activation^12^. Changes in cytoplasmic calcium concentration thus could provide a mechanistic link between BCR signal strength, the switch-on of downstream effector processes and their temporal regulation.

In contrast to qualitative description, absolute quantification of cytosolic calcium has not been achieved yet, partly because of the lack of internal concentration standards. Two-fluorophore Förster resonance energy transfer (FRET)-GECI, that can take on a calcium-saturated (quenched) and calcium-unsaturated (unquenched) condition, could overcome this, however, its intravital application has been hampered by light distortion effects in deeper tissue. The differential scattering and photobleaching properties of the two fluorophores would lead to a false bias towards a higher quenching state. We here introduce a single-cell fluorescence lifetime imaging (FLIM) approach for absolute calcium quantification in living organisms that is tissue depth-independent. The eCFP/citrine-FRET pair-GECI TN-XXL is able to measure fluctuations in cytoplasmic calcium concentration through the calcium binding property of the muscle protein Troponin C (TnC)^13^. Calcium binding to the fluorophore-linker TnC quenches eCFP fluorescence through energy scavenging by citrine, linking decreasing eCFP fluorescence lifetime to increasing calcium concentration. In addition, phasor analysis of FLIM data elegantly condenses multicomponent fluorescent decay curves into single vector-based information (the phasor)^14^. For calcium concentration analysis in microscopic images, we took advantage of this by projecting the phasor value in each pixel onto a given calibration^15^. With this method, we are able to describe short- and long-term changes in absolute calcium concentrations within B cells during affinity maturation and differentiation into antibody-producing plasma cells.

We here show, using our calcium reporter mouse strain “YellowCaB” (<u>yellow</u> fluorescence after <u>ca</u>lcium influx in <u>B</u> cells), which expresses cytosolic TN-XXL under control of the CD19 promoter, that calcium signals are highly dynamic within different B cell populations. We analyze AG-specific and non-AG-specific extrafollicular and GC B cells as well as AG-specific extrafollicular plasma blasts. We describe highly dynamic signaling patterns that differ in amplitude and baseline and correlate with cellular differentiation stages.

## Results

### YellowCaB: A system for FRET-based calcium analysis in B cells

By breeding mice expressing a loxP-flanked STOP sequence followed by the TN-XXL-construct inserted into the ROSA26 locus together with the CD19-Cre strain^16^, we generated offspring with exclusive and visible expression of the FRET fluorophore pair eCFP and citrine in CD19^+^ B lymphocytes, as confirmed by confocal microscopy after magnetic B cell isolation (Fig 1a and b). YellowCaB cells were excited with a 405nm Laser that is capable of exciting eCFP but not citrine. The detection of yellow emission thus can be attributed to baseline FRET representing steady state calcium levels. Expression of TN-XXL in YellowCaB mice was tested by flow cytometry, determining the percentage of fluorescent cells in a CD19^+^GFP^+^ gate after excitation with the 488nm laser. Yellow citrine fluorescence was found to be exclusive to CD19^+^ B lymphocytes and the expression level of TN-XXL ranged between 25-45% in mice expressing the sensor on one allele (CD19^cre/+^ TN-XXL^+/-^) and 70-90% in mice homozygous for the TN-XXL construct (CD19^cre/+^ TN-XXL^+/+^) animals (Fig 1c and S1). When analyzing secondary lymphoid organs, no differences in total cell numbers and B cell numbers between TN-XXL^+/-^ CD19^cre/+^, TN-XXL^+/+^ CD19^cre/+^ and wild type mice were detected (Fig S1). We next set out to test if we could induce a FRET signal change under calcium-saturating conditions in the cytoplasm. The ionophore ionomycin is commonly used as positive control in *in vitro* experiments measuring calcium concentrations, as it uncouples the increase of calcium concentration from the physiological entry sites of Ca^2+^ ions by forming holes in the cell membrane. When stimulated with ionomycin, a steep increase of the FRET level over baseline could be recorded by flow cytometry in the GFP-channel after excitation with the 405nm laser. Calcium-dependence was independently confirmed by staining with the calcium sensitive dye X-Rhod-1 that shows a red fluorescence signal increase after calcium binding (Fig 1d)^17^.

**Fig. 1.**
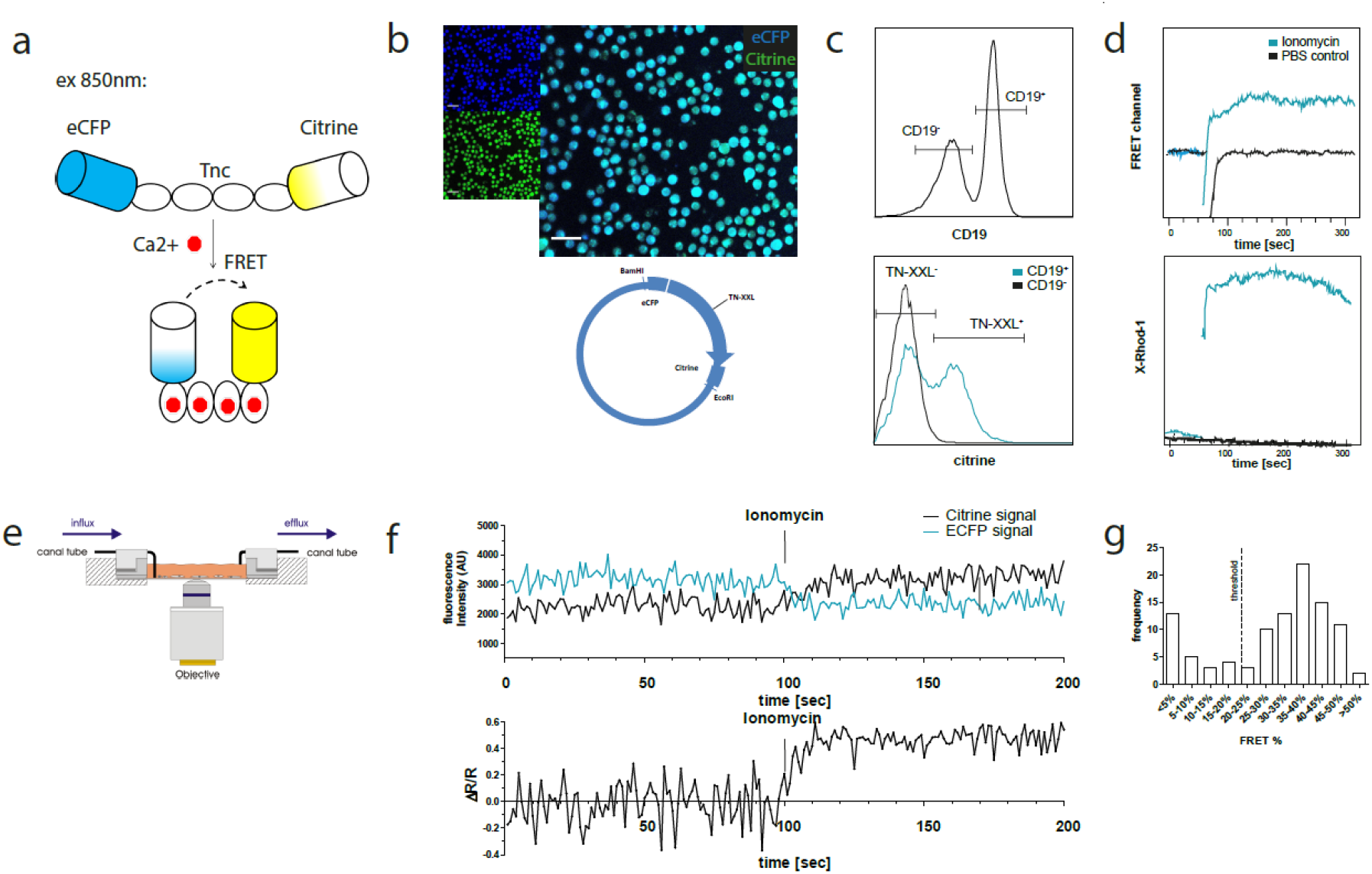
The GECI TN-XXL is functionally expressed in CD19^+^ B cells of YellowCaB mice. a Schematic representation of the genetically encoded calcium indicator TN-XXL with calcium sensitive domain TnC fused to donor fluorophore eCFP and acceptor fluorophore citrine. Binding of Ca^2+^ ions within (up to) four loops of TnC leads to quenching of eCFP and Förster resonance energy transfer to citrine. b Confocal image of freshly isolated CD19^+^ B cells. Overlapping blue and yellow-green fluorescence of eCFP and citrine, respectively, can be detected after cre-loxP mediated expression of the TN-XXL vector in YellowCaB mice. c Flow cytometric analysis of TN-XXL expression among lymphocytes of YellowCaB mice. d Flow cytometric measurement of calcium flux after addition of ionomycin and PBS control. e Continuous perfusion imaging chamber for live cell imaging. f Confocal measurement of mean fluorescence intensity and FRET signal change after addition of ionomycin to continuous perfused YellowCaB cells. Data representative for at least 100 cells out of three independent experiments. g Frequency histogram of >100 YellowCaB single cells, FRET analyzed after ionomycin stimulation. Threshold chosen for positive FRET signal change = 20% over baseline intensity.

In preparation of our intravital imaging experiments, we first needed to test if the YellowCaB system is stable enough for time-resolved microscopic measurements and sensitive enough for subtle cytoplasmic calcium concentration changes as they occur after store-operated calcium entry (SOCE). In SOCE, stimulation of the BCR with AG leads to drainage of intracellular calcium stores in the endoplasmic reticulum (ER) which triggers calcium influx from the extracellular space into the cytosol through specialized channels^18^. We established a customizable continuous perfusion flow chamber image system to monitor and manipulate YellowCaB cells over the duration of minutes to hours (Fig. 1e). Division of the fluorescence intensity of electron acceptor citrine by that of donor eCFP yields the FRET ratio (R), which is then put into relationship to baseline FRET levels. As citrine intensity is expected to increase and eCFP intensity is expected to decrease due to FRET, the resulting value of ΔR/R will increase as well. As expected, we could detect a decrease of the CFP signal, concurrent with an increased YFP fluorescence after the addition of 4μg/ml ionomycin to continuous flow of 6mM Krebs-Ringer solution. Overall, this resulted in a maximal elevation of ΔR/R of 50-55% over baseline (Fig. 1f). Analysis of >100 cells showed that approximately in three quarters of the cells we were able to detect FRET in response to ionomycin treatment, and that the majority of these cells showed 35-40% FRET signal change. According to the two populations visible in the histogram, we defined a change of 20% ΔR/R as a relevant threshold for the positive evaluation of responsiveness (Fig. 1g). In conclusion, we achieve the functional and well tolerated expression of TN-XXL exclusively in murine CD19^+^ B cells for measurement of changes of cytoplasmic calcium concentrations. Moreover, flow cytometric and microscopic long-term calcium analysis are possible.

### Repeated BCR stimulation results in fluctuating cytoplasmic calcium concentrations

SOCE in B cells can be provoked experimentally by stimulation of the BCR with multivalent AG, for example anti-Ig heavy chain F(ab)_2_ fragments. To test the functional performance of the GECI TN-XXL in YellowCaB cells, we stimulated isolated YellowCaB cells with 10 μg/ml anti-IgM F(ab)_2_ fragments to activate the BCR. In an open culture imaging chamber, we could induce an elevated FRET signal with a peak height of >30% that lasted over three minutes (Fig 2a). The signal declines after this time span, probably due to BCR internalization or the activity of ion pumps. We tested antibody concentrations at 2 μg/ml, 4 μg/ml, 10 μg/ml and 20 μg/ml. An antibody concentration of 2 μg/ml was not enough to provoke calcium flux (data not shown), whereas at 4 μg/ml anti-IgM-F(ab)_2_ we could observe 20% elevated ΔR/R over baseline (Fig. 2b). At 20 μg/ml anti-IgM-F(ab)_2_ we could see no further FRET increase (Fig S2). Thus, we conclude a concentration dependency of the GECI TN-XXL and saturating conditions at 10 μg/ml BCR heavy chain stimulation. Interestingly, the reaction is not completely cut off after the FRET signal has declined, but a residual FRET signal of about 7% compared to baseline values can be measured for approximately 3.5 additional minutes (Fig. 2a). Thus, B cells seem to be able to store extra calcium within the cytoplasm for some time. We therefore asked if it is possible to stimulate YellowCaB cells more than once. Indeed, we could stimulate YellowCaB cells *in vitro* repeatedly with F(ab)_2_-fragments of anti-IgM-antibody. For this purpose, we connected our imaging culture chamber to a peristaltic pump and took advantage of the fact that under continuous perfusion with Ringer solution, the flow will dilute the antibody out of the chamber. This way, it is possible to stimulate B cells several times rapidly and subsequently, before BCRs are internalized (Fig. 2b), indicated by multiple peaks in ΔR/R. We repeated this procedure several times and could observe this type of repetitive response up to five times. Also, stimulation of the BCR light chain using an anti-kappa antibody leads to calcium increase within YellowCaB cells (Fig. S2). Of note, the resulting FRET peak is shaped differently, and concentrations >150μg/ml antibody were needed in order to generate a response. This might be in part due to the fact that monovalent AG is not sufficient to drive BCR activation and the light chain has different conformational properties.

**Fig. 2.**
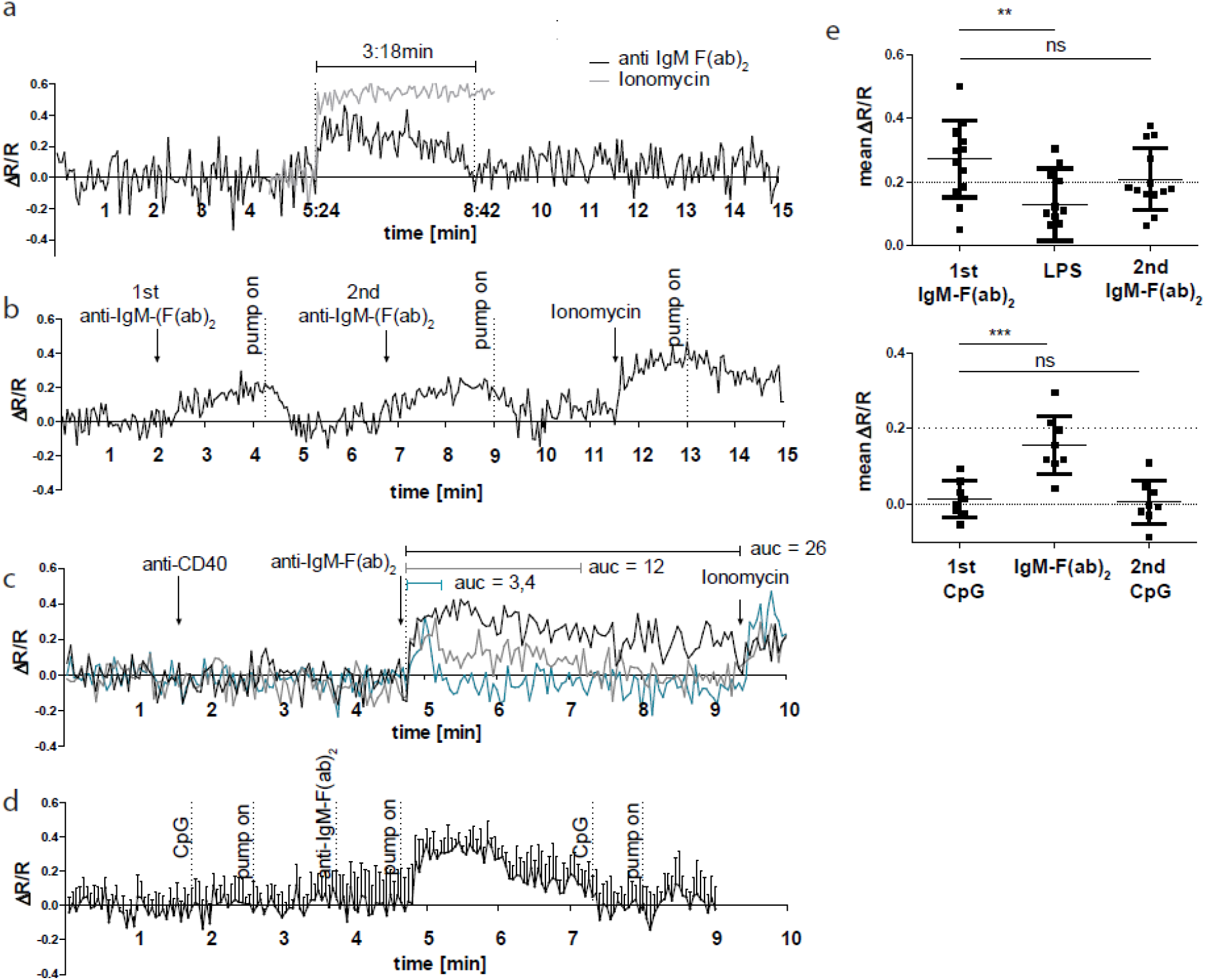
BCR stimulation specifically leads to calcium mobilization in YellowCaB cells *in vitro.* a Confocal measurement of FRET duration (ΔR/R>0) in non-perfused primary polyclonal YellowCaB cells after addition 10μg/ml anti-IgM-F(ab)_2_ (black), Ionomycin control (grey). Data representative for at least 35 single cells in four independent experiments. b Confocal measurement of FRET signal change after repeated addition of anti-IgM-F(ab)_2_ to perfused primary polyclonal YellowCaB cells. Data representative for at least 50 cells out of five independent experiments. c Confocal measurement of FRET signal change after addition of anti-IgM-F(ab)_2_ to perfused primary polyclonal YellowCaB cells following stimulation with anti-CD40 antibody and Ionomycin positive control. Examples of transient cytoplasmic (blue), intermediate (grey) and sustained calcium mobilization shown, area under the curve (AUC) compared. Data representative for 26 cells out of two independent experiments. d Resulting FRET curve out for n=7 primary polyclonal YellowCaB cells perfused with TLR9 stimulator CpG in Ringer solution and subsequent addition of anti-IgM-F(ab)_2_. e Mean FRET signal change over time after addition of TLR4 or TLR9 stimulation in combination with BCR crosslinking by anti-IgM-F(ab)_2_ in perfused polyclonal YellowCaB cells n=12 (upper) and n=8 (lower) **: p=0.0086, ***: p<0.05, one-way ANOVA. Error bars: SD/mean.

Since T cell engagement and the binding of unspecific microbial targets to innate receptors like Toll like receptors (TLRs) have also been described to raise cytoplasmic calcium in B cells^19–21^, we investigated the response of YellowCaB cells after incubation with anti-CD40 antibodies, as well as the TLR4 and TLR9 stimuli Lipopolysaccharide and cytosine-phosphate-guanin-rich regions of bacterial DNA (CpG), respectively. Within the same cells, we could detect no reaction to anti-CD40 treatment alone, but observed three types of shapes in post-CD40 BCR-stimulated calcium responses, that differed from anti-CD40-untreated cells (Fig. 2a). These calcium flux patterns were either sustained, transient or of an intermediate shape (Fig. 2c). Sustained calcium flux even saturated the sensor at a level comparable to that achieved by ionomycin treatment. Cells that showed only intermediate flux maintained their ability to respond to ionomycin treatment at high FRET levels, as demonstrated by the ΔR/R reaching 0.4 again after stimulation (Fig. 2c). Furthermore, integrated TLR and BCR stimulation affected the appearance of the calcium signal. The addition of TLR9 stimulus CpG alone had no effect on YellowCaB FRET levels, however, the subsequent FRET peak in response to anti-Ig-F(ab)_2_ was delayed (Fig 2d+e). TLR4 stimulation via LPS could elevate calcium concentration of B cells, but only to a minor extent (Fig 2e). Pre-BCR TLR4 stimulation by LPS lead to decreased FRET levels in response to anti-IgM-F(ab)_2_. We conclude that, in order to get fully activated, B cells are able to collect and integrate multiple BCR-induced calcium signals and that signaling patterns are further shaped by innate signals or T cell help. BCR-inhibition abolishes a FRET signal change in response to anti-IgM-F(ab)_2_ (Fig S2). Of note, we excluded the possibility that measured signal changes were related to chemokine stimulation. *In vitro,* we could detect no FRET peak after applying CXCL12, probably because of lacking GECI sensitivity to small cytoplasmic changes (Fig. S2). Thus, the YellowCaB system is well suited for the detection of BCR-induced cytosolic calcium concentration changes.

### Fluctuating calcium levels are observed *in vivo*

We next set out to investigate if the ability of B cells to collect calcium signals is also shared by germinal center B cells, and to identify the spatial distribution of possible BCR-triggering forces in an *in vivo* setting. For two-photon intravital imaging, nitrophenyl (NP)-specific B1-8hi:YellowCaB cells were transferred into wild type hosts which were subsequently immunized with NP-CGG (chicken gamma globulin) into the right foot pad^22^. Mice were imaged at day 8 p.i. when GCs had been fully established. Activated TN-XXL^+^ YellowCaB cells had migrated into the GC and, as confirmed by positive PNA- and anti-GFP immunofluorescence histology (Fig 3 a). At this time point, mice were surgically prepared for imaging as described before^23^. Briefly, the right popliteal lymph node was exposed, moisturized and flattened under a cover slip sealed against liquid drainage by an insulating compound. The temperature of the lymph node was adjusted to 37°C and monitored during the measurement. Our experiments revealed that the movement of single YellowCaB cells can be tracked *in vivo. C*alcium fluctuations can be made visible by intensity changes in an extra channel that depicts the FRET signal, as calculated from relative quenching of TN-XXL. Color-coding of intensity changes in the FRET channel showed time-dependent fluctuations of the signal and, in some particular cases, a sustained increase after prolonged contacts between two YellowCaB cells (Fig 3 b, movie S1). Interestingly, FRET intensity seemed to be mostly fluctuating around low levels in moving cells, whereas sustained increase required cell arrest (Fig S3). We already showed that signal changes in FRET of TN-XXL are reflecting BCR activation. The observed calcium fluctuations might therefore coincide with cell-to-cell contacts between FDCs and B cells, resulting in AG-dependent BCR stimulation. To test for this, we first measured the colocalization between signals within the FDC-channel and the citrine channel. The intensity of colocalization I_coloc_ of all cells was plotted as a function of frequency and biexponentially fitted (Fig 3 c). We set the threshold for a strong and sustained colocalization of FDCs and B cells to an intensity of 150 AU within the colocalization channel. At this value, the decay of the biexponential fit was below 10%. We thus decided to term all cells with a colocalization intensity = 0 (naturally the most abundant ones) not colocalized, cells with a colocalization intensity between 1 and 150 transiently colocalized to FDCs (“scanning” or shortly touching the FDCs) and all cells above this intensity threshold strongly or stably colocalized. When we compared the relative FRET intensity changes ΔR/R of a tracked cell (Fig 3 b, cell 1), where baseline R is the lowest FRET intensity measured, and its contacts to FDCs, we could indeed detect several transient B-cell-FDC contacts that were followed by a step-wise increase of ΔR/R and thus an increase of cytoplasmic calcium concentration (Fig 3 d). These increases in B cells were not only restricted to contacts with FDCs, but also occurred between B cells: A visible contact of cell 1 (Fig 3 b, cell 1) to a fellow B cell (Fig 3 b, cell 2) caused a sustained boost of the calcium concentration in the tracked cell 1 (Fig 3d). Cell 2 itself kept strong FDC contact over the whole imaging period and maintained elevated, but mostly stable ΔR/R values. These experiments confirmed that GC B cells are able to collect calcium as a consequence of repeated signaling events mediated by FDC-to-B cell contacts and, surprisingly, also by B-to-B cell contacts *in vivo*.

**Fig. 3.**
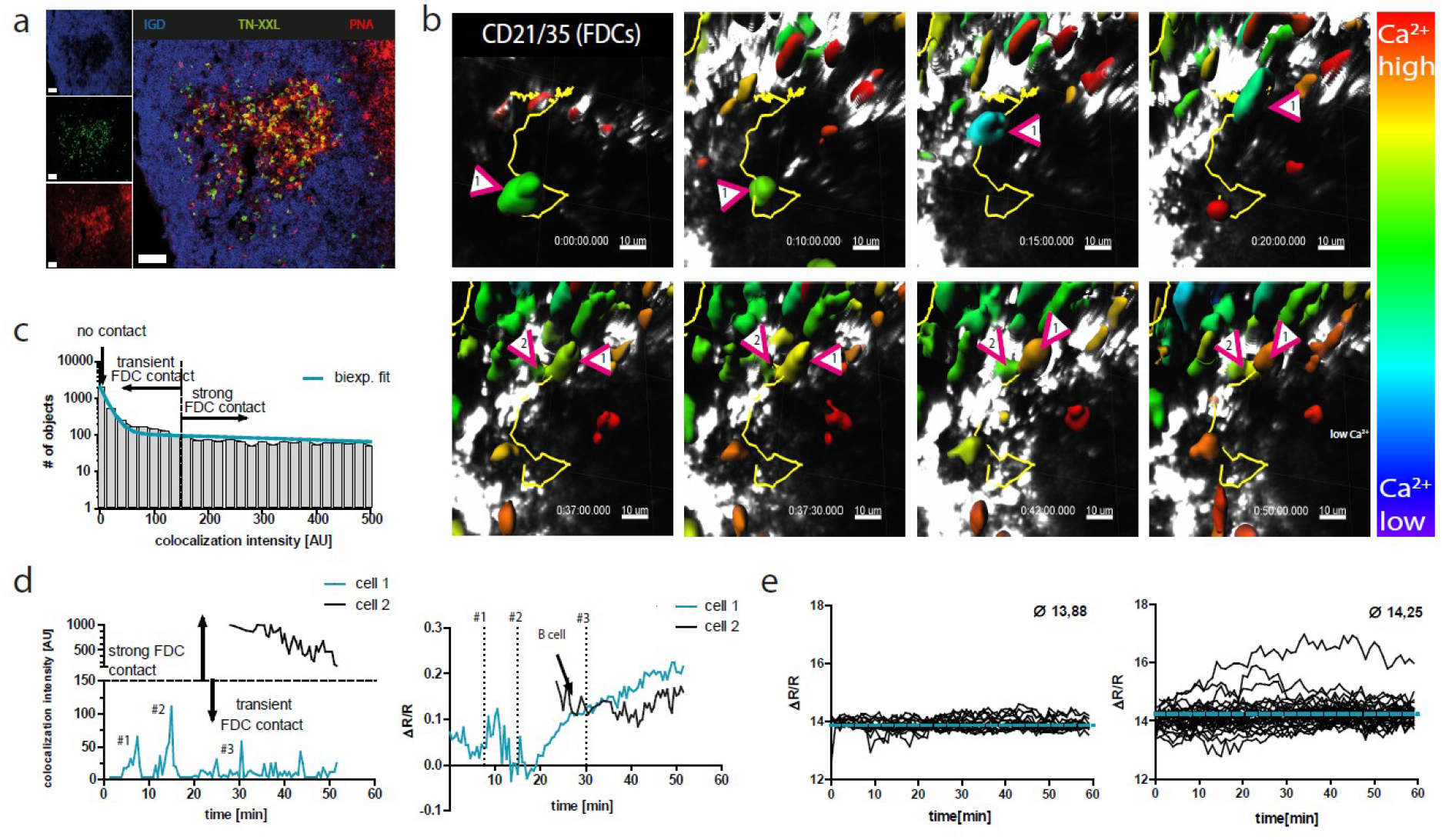
YellowCaB cells form productive germinal centers in vivo and show active BCR signaling after cell-to-cell contacts. a Histological analysis of lymph nodes after adoptive transfer of polyclonal YellowCaB cells to restricted hosts. TN-XXL (green) positive cells cluster in IgD (blue) negative regions and are showing activation, confirmed by PNA staining (red). Scale bar 50μm. b Stills of intravital imaging of polyclonal YellowCaB cells transferred to restricted host. 3D surface rendering and single-cell tracking (track line in yellow) with relative color coding ranging from blue= low ΔR/R to red=high ΔR/R c Histogram showing cell frequency vs. colocalization intensity [AU] and biexponential fit of data. A curve decay of <10% was set as threshold parting transient from strong B cell-to-FDC contact. All cells with colocalization intensity <1 were assigned negative. d Colocalization intensities of tracked cells 1 and 2 over time vs FRET signal change of cell 1 and 2 over time. Contact events to FDCs are assigned numbers #1, #2 and #3. Contact between cell 1 and 2 is indicated by arrow. e Comparison of FRET signal change of naive and AG-specific YellowCaB cells over time.; blue line indicates mean value of all tracked cells n=16 (naive) and n=33 (AG-spec.).

Next, we asked if the ability to perform BCR signaling is dependent on BCR affinity. Thus, we adoptively transferred stained polyclonal, non-AG-specific YellowCaB cells one day prior to intravital imaging and compared FRET-signal changes of several tracked cells over time within the same measurement. Non-AG-specific YellowCaB cells showed a rather homogenous distribution of calcium concentration with low intensity fluctuations in the FRET channel around a mean of 13.88 AU, whereas AG-specific YellowCaB cells showed heterogeneous signaling patterns with higher FRET intensities of 14.25 AU on average (Fig. 3e). We compared ratiometric FRET histograms of non-AG-specific and AG-specific cells out of five different measurements. To do so, we normalized FRET values by the mean fluorescence intensity averaged over all non-AG-specific YellowCaB cells. This further confirmed a positive correlation of B cell calcium concentration and BCR-affinity (Fig S4).

### Absolute quantification of cytoplasmic calcium by eCFP fluorescence lifetime analysis reveals activation-dependent calcium heterogeneity in B cells

Due to varying imaging depths and therefore differing noise levels in tissue, the comparison of measurements in a ratiometric set-up only gives relative information about calcium concentrations. Therefore, universal statements on calcium levels among different B cell populations will demand absolute quantification. We switched our imaging set-up from the analysis of the fluorescent intensities of eCFP and citrine to the measurement of the fluorescence lifetime (tau) of the FRET-donor eCFP. Fluorescence lifetime is defined as the mean time a fluorescent molecule stays in an elevated energetic state after excitation, before photon emission and relaxation to the ground state take place. As a fully calcium-quenched eCFP in the GECI TN-XXL would transfer its energy mainly to citrine, its fluorescence lifetime would be measurably shorter than on an unquenched eCFP. Phasor analysis of time-domain FLIM virtually transfers time-resolved fluorescence data into phase domain data by discrete Fourier transformation^14^. This approach overcomes the obstacles of multi-component exponential analysis and yields readily comparable pixel-or cell based plots that assign a position within a half-circle to each data point, dependent on the mixture of lifetime components present^24^.

Employing our adoptive cell transfer set up described above (Fig 4 a), we could divide GC B cells into five different populations based on their location in the imaging volume and their fluorescent appearance. At day 8 p.i., polyclonal YellowCaB cells, identified by their red labeling, mostly lined up at the follicular mantle around the GC, with some of them having already entered into activated B cell follicles. AG-specific, citrine-positive B1-8hi:YellowCaB cells were found clustered in the GC, close to FDCs, outside of GC boundaries, or as bigger, egg-shaped differentiated cells in the extrafollicular medullary cords (MC), probably comprising plasma blasts (Fig. 4 b). Color-coded 2D and 3D FLIM analysis of these populations confirmed that calcium concentrations were fluctuating within all of those B cell populations, and that most B cells were maintaining relatively high fluorescence lifetimes and therefore low calcium concentration on average with single-cell exceptions (Fig. 4 b, movie S2). It should be noted that autofluorescence of the capsule or macrophages contributed to tau values <800ps, indicated by dark red color-coding and is not to be attributed to high calcium values. Phasor analysis and plotting of the B cell-wise segmented lifetime data further confirmed the presence of cell clones with quenched TN-XXL, somewhat surprisingly also among plasma blasts, which are thought to down-regulate their surface BCR (Fig 4c, movie S3). To translate the measured lifetime values into absolute calcium concentrations, we projected the data points onto the phasor connecting quenched and unquenched eCFP (Fig. 4d). In this way, we corrected for artifacts acting on the phasor vectors and, implicitly, on the fluorescence lifetimes caused by contribution of background noise (Fig S5). Comparison of AG-specific cells inside GCs with those outside GCs, and non-AG-specific cells inside GCs as well with those outside GCs showed that the distribution of calcium concentrations of these B cells were dependent on BCR specificity and rather independent from their location within the imaging volume, despite higher fluctuation seen among AG-specific populations (Fig S5). However, we noted the emergence of a cell subset that is high in calcium and therefore located on the right half of the plot, in AG-specific cells and most prominent among cells within the MC, as compared to non-AG-specific YellowCaB cells. Overlay of an imaging snapshot shows that only few cells are in a state of elevated calcium (>800 nM) at one given time point (Fig. 4e). Accordingly, these maxima were reached as transient fluctuation peaks, i.e. periods of below one minute, in which these concentrations seem to be tolerated. Calcium values exceeding the dynamic range of TN-XXL (>857nM) were recorded for all measured subsets, but the most cells >857nM were found among intrafollicular AG-specific B cells and extrafollicular AG-specific B cells. (Fig. 4f). The heterogeneity in temporal calcium concentrations therefore is smallest among non-AG specific B cells, increases with activation in AG-specific GC B cells, and is most prominent among plasma blasts. Thus, a progressive heterogeneity of calcium signals within B cells can be seen alongside the process of activation and differentiation.

**Fig. 4.**
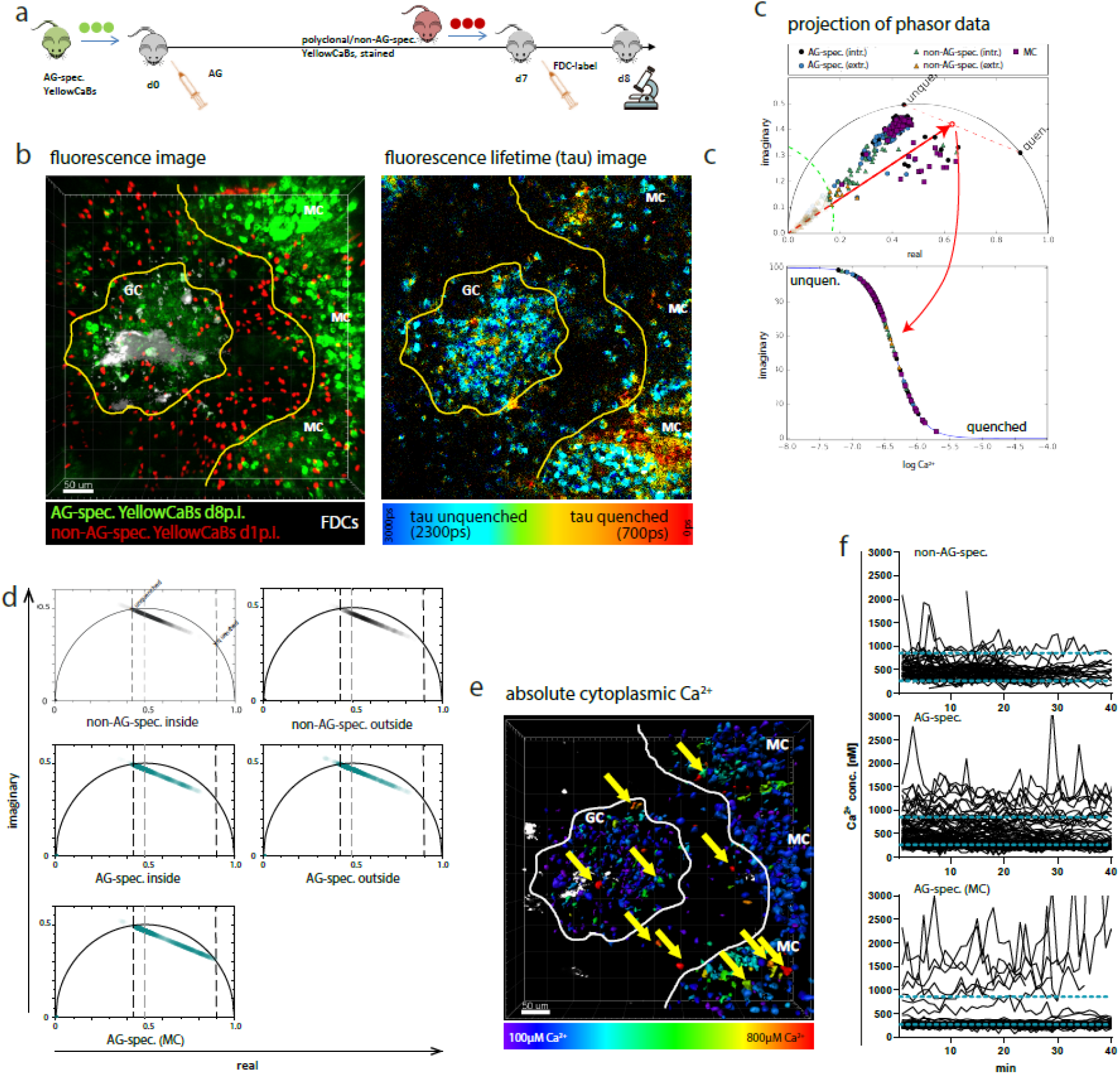
Determination of absolute calcium concentration by intravital fluorescence lifetime imaging of GC B cell populations. a Cell transfer and immunization strategy for intravital imaging of AG-specific and polyclonal YellowCaB B cells. b z-stack of intravitally imaged germinal center (GC) and medullary cords (MC). B cells have been distinguished into polyclonal, non-AG-specific YellowCaB cells (red) inside and outside the germinal center, AG-specific YellowCaB cells inside and outside the germinal center and differentiated B cells (plasma blasts) due to staining and localization (left). Color-coded fluorescence lifetime image (right) with lifetimes of unquenched eCFP depicted in blue and lifetimes of quenched eCFP in red. c Phasor plot of fluorescence lifetimes measured in b with indicated subpopulations. Green radius indicates noise that has been excluded from the evaluation of absolute calcium concentrations. For absolute calcium concentration determination, phasors of single cells have been projected to the red-dashed line connecting fully calcium-saturated (quenched) and calcium-free (unquenched) conditions d Projections of phasor vectors onto connecting line between unquenched and quenched eCFP lifetimes among the different subpopulations. f calcium concentrations intravitally measured with FLIM-phasor in germinal center B cell populations over time. Non-AG-specific YellowCaB cells (left, n=92) compared to AG-specific YellowCaB cells (middle, n=169) and extrafollicular, AG-specific YellowCaB cells (right, n=69) The dynamic range of the GECI TN-XXL is indicated by blue dashed lines.

### Functional relevance of increased calcium concentration among extrafollicular YellowCaB cells

The surprising finding of elevated cytoplasmic calcium levels in extrafollicar B cells led us to analyze this subset further. Recently, contacts with subcapsular sinus macrophages (SCSM) were reported to induce the differentiation of memory B cells to antibody producing plasma cells^25–27^. To test, if SCSM could be the cause of elevated extrafollicular calcium levels in differentiated, extrafollicular B cells, we. intravitally imaged wild type host mice that have been adoptively transferred with B1-8hi:YellowCaB cells and received an injection of efluor660-labeled anti-CD169 antibody together with the usual FDCs labeling one day prior to analysis. We concentrated on the area beneath the capsule, identified by second harmonic generation signals of collagen fibers in this area. Thresholds of colocalization between CD169^+^ macrophages and TN-XXL^+^ YellowCaB cells are described in Fig S6. Together, these methods led to the 3D visualization of the SCS with CD169 stained macrophages, lined up in close proximity. (Fig 5a). AG-specific YellowCaB cells were detected clustering in GCs nearby. Extrafollicular YellowCaB cells crowding the SCS space were found to have multiple contact sites to SCSM. Some B cells were observed to migrate along the SCS, possibly scanning for stimulatory signals like antigen (movie S4). Cell tracking and simultaneous analysis of absolute calcium concentration and colocalization intensity revealed that B-cell-to-SCSM contacts resulted in an immediate increase in cytoplasmic calcium concentration. In line with this, the loss of a transient-made contact directly leads to a decrease in calcium concentration (Fig 5b). This was further confirmed by bulk analysis of all detected YellowCaB cells over the whole imaging period (Fig 5c). Relating calcium concentration and colocalization showed that the calcium concentration in YellowCaB cells with direct contact to SCSM reaches values that are more than doubled compared to that in cells that were not in contact Furthermore, calcium concentration increase seems to be positively correlated to B cell-to-SCSM contact strength. Colocalization intensities between 0 and 1 were defined as no-contact. All values above describe transient or strong contact. The threshold between transient and strong contact has been set to 10% decay of the biexponentially fitted histogram over all cells. Cells that were defined as strong contacters also reached the highest calcium concentrations. We conclude that contacts of B cells to SCSM could induce elevation of B cell cytoplasmic calcium concentrations, presumably due to activation, with the absolute concentrations being dependent on the strength of the contact.

**Fig. 5.**
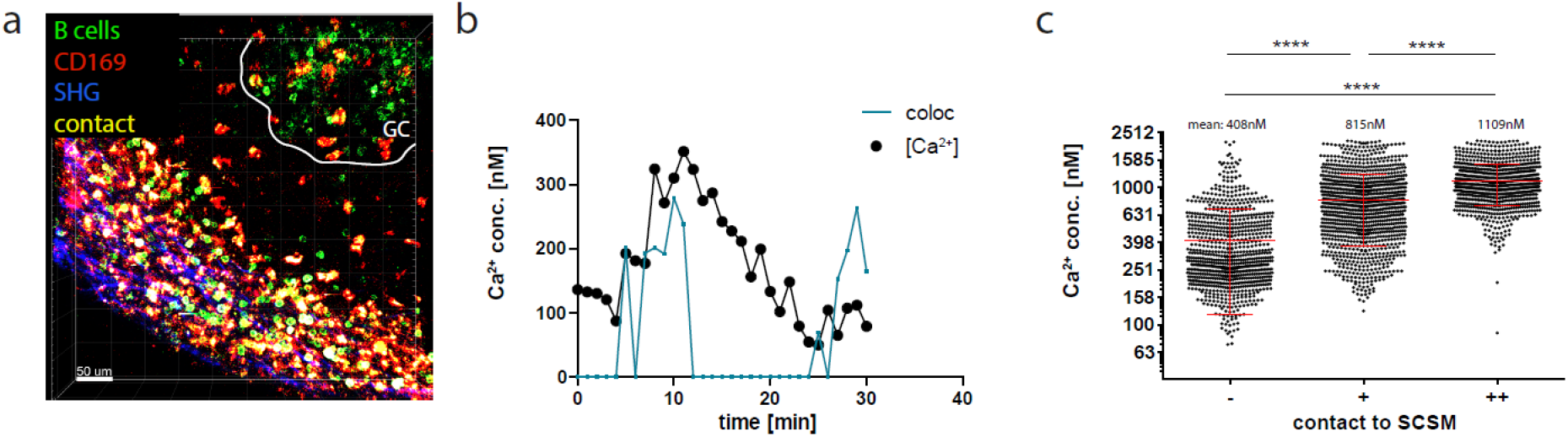
Differentiated B cells show elevated calcium concentrations after contact to macrophages of the subcapsular sinus. a z-stack of intravitally imaged lymph node with germinal center (GC) and subcapsular sinus (SHG, blue). CD169^+^ macrophages (red) lining the subcapsular sinus show large contacts (yellow) to YellowCaB cells (green) of increased brightness and size that therefore have been identified as differentiated B cells. Size 500×500×78μm. Scale bar 60μm. b Single-cell track of a YellowCaB cell making transient contact to macrophage, blue: colocalization intensity [AU], black: change of absolute calcium concentration. c FLIM measurement of mean absolute calcium concentration of YellowCaB cells showing no (-), transient (+) or strong (++) overlap with CD169^+^ signal. n=1000, ANOVA analysis (****: p<0.0001), mean and SD.

## Discussion

Intravital imaging technologies have helped to elucidate the nature of the GC reaction a great deal. AG-capture, cycling between zones, and development of clonality patterns have been made visible by two-photon microscopic techniques^5,28–30^. Furthermore, important functional *in vivo* data like signaling in T helper cells have been collected using a calcium sensitive protein^31,32^. However, research on signaling of (GC) B cells *in vivo* so far involved transfer and immunization experiments and *ex vivo* analysis of sorted cells, or used non-reversible BCR-signaling reporters like Nur77, leaving aside the notion that B cell activation and selection is a highly dynamic process^10,33,34^. These experiments concluded that only a small amount of GC B cells seem to maintain the signaling capacity of naive, mature B cells and that BCR signaling thus is dispensable in GCs^10,11,35–38^. On the other hand, BCR-regulating surface proteins like CD22 or Sialic acid-binding immunoglobulin-type lectins (Siglecs) have been related to development of autoimmunity and point out to BCR signaling as an important component, not only in B cell development but also in differentiation to effector cells^39–42^.

For flexible analysis of signaling in cells of the CD19^+^ lineage, we developed of a novel transgenic reporter system and image processing approach, enabling quantification of the signaling second messenger calcium. For the first time, we could measure absolute B cell cytoplasmic calcium concentrations *in vivo.* The FRET-based GECI TN-XXL can be used stably in moving, proliferating and differentiating lymphocytes and the reversibility of the sensor makes it suitable for longitudinal intravital measurements.

At first, by measuring calcium flux within B cells *in vitro*, we could show fast reversibility and suitable sensitivity of the GECI TN-XXL for intravital applications. Using a flow chamber system, we could show that repeated BCR activation leads to repeated calcium elevation in the cytoplasm. Thus, our system shows that B cells are able to collect and integrate sequential signals, probably for positive selection, acting via calcium accumulation up to a hypothetic threshold. In support of that, B cellular calcium concentration must not constitutively exceed a certain value in order to prevent mitochondrial depolarization^43,44^ On the other hand, it is known that certain elevated calcium concentrations and timely regulated calcium flux patterns decide if and which downstream transcription factors are activated^45,46^. Balancing out calcium concentrations might thus also function in sensing the completion of affinity maturation. We propose calcium fluctuation as time- and strength-coded system for the transduction of information about growing BCR affinity. Other fine-tuned signaling mechanisms are well known to instruct transcriptional mechanisms, though partly conflicting; e.g. CD40 signaling is absolutely indispensable for entry of B cells into the GC reaction, but prolonged CD40 signaling instead promotes IRF4 transcription and therefore favors an extrafollicular ASC phenotype^47^.

Next, to investigate B cell signaling using an intravital set-up, we needed to establish an intensity- and scattering-independent approach. Using phasor-based FLIM, we were able to master the hindering effects of wavelength-dependent aberrations in the tissue that influence the performance of each fluorophore individually. In order to extract absolute cytoplasmic calcium concentrations from the donor lifetime data *in vivo,* we developed a numerical strategy based on the exclusion of low SNR pixel values and on the correlation of amplitude and phase of the phase vectors to account for experimental uncertainty.

Employing this methodology, we can describe the connection between signaling and selection in a novel way. We could visualize that during differentiation, B cells undergo repeated stimulation via BCR- and costimulatory signals that stem from serial contacts of B cells with FDCs or other B cells, the latter possibly being explained by simultaneous stimulation of BCRs and Fc receptors^48–52^. Using time-resolved FRET-FLIM measurements in GC, we could show that BCR AG-specificity and state of differentiation are closely related to distinct degrees of heterogeneity of calcium concentrations. Furthermore, we found that differentiated extrafollicular B cells or plasma blasts are among the cells with highest calcium concentrations within our set up. It is known that high calcium concentrations might also act as stress signal that demands restriction. Recent reports state that elevated calcium might set a “metabolic timer” for T cell help to rescue a mitochondria-depolarizing ion concentration^53^. This shows that calcium levels within cells need to be tightly regulated, as it does also play its part in the induction of apoptosis. Calcium levels of >1 μM over the duration of >1 h have been reported damaging in neurons^54,55^. Since apoptosis is the default fate for B cells in the germinal center reaction^56^, CD40 and TLR signaling might contribute to the limiting of cytoplasmic calcium concentrations and thus promote cell survival of B cell clones with appropriate BCR affinity^21,57–60^. For CD40 signaling in immature B cells this has been confirmed^61^. Our data does show that TLR signaling can attenuate calcium flux in stimulated B cells, CD40 can either attenuate or augment the calcium response, presumably depending on the affinity of the BCR and its efficiency in presenting AG^29,32^.

We are confident that our set-up models different B cell activation and differentiation stages in a single measurement, though it is worthwhile noting that the cells imaged within the MC region most likely comprise of extrafollicularly generated plasma blasts, rather than differentiated B cells that have left the GC (these are reported to appear only at much later stages of the response)^62^. We suggest that during differentiation, calcium signals mainly act as transcriptional regulators, whereas it is likely that this is different in terminally differentiated cells. Indeed we were able to image ongoing cytoplasmic calcium elevation in the regions of the MC or the subcapsular sinus and confirmed that B cells in contact to SCSM had significantly higher cytosolic calcium concentrations. The SCS has recently been proposed as site of reactivation of memory B cells^25^.

Importantly, changes in mitochondrial membrane potential and/or the integrity of the ER also lead to varying calcium concentrations within cells, since both act as major intracellular calcium buffering organelles^63^. A close connection between mitochondrial calcium homeostasis, altered ROS production and the expression of plasma cell master transcription factor BLIMP1, as well as changes in metabolism have been reported previously^64,65^. We have already applied phasor-FLIM of endogenous NAD(P)H fluorescence for mapping of enzyme activities in cell cultures^62^. The application of this method and combination with calcium imaging holds great potential to further dissect immunometabolic processes in B cells, short- and long-lived plasma cells *in vivo*.

## Methods

### Mice

PR26CAGTN-XXL^flox/flox^ mice were obtained by courtesy of F. Kirchhoff, Saarland University, Homburg^66^. YellowCaB mice were generated by crossing of R26CAGTN-XXL^flox/flox^ mice with the CD19^cre/cre^ strain^16^ and maintained on a C57/Bl6 background. Only YellowCaB mice heterozygous for cre were used to avoid deletion of CD19. Mice with monoclonal NP-specific BCR (B1-8^hi^:YellowCaB) were generated by crossing of YellowCaB mice with B1-8^hi^ mice^22^. Thx mice were used as irrelevant-BCR hosts. All mice were bred in the animal facility of the DRFZ. All animal experiments were approved by Landesamt für Gesundheit und Soziales, Berlin, Germany, in accordance with institutional, state, and federal guidelines.

### Cells

Primary splenocytes were isolated from whole spleens of YellowCaB mice or B1-8^hi^:YellowCaB mice in 1xPBS and erythrocytes lysed. B cells were negatively isolated using the Miltenyi murine B cell isolation kit via magnetic assisted cell sorting (MACS) leave B cells untouched in order not to pre-stimulate them.

### Staining and flow cytometry

Single cell suspensions were prepared and stained according to the guidelines for flow cytometry and cell sorting in immunological studies^67^. To simultaneously assess calcium influx with a dye-based method we stained whole splenocytes or isolated B cells with the calcium sensitive dye X-Rhod-1 (invitrogen). X-Rhod-1 is a single-fluorophore calcium reporter molecule that enhances its fluorescence intensity upon calcium binding in a range of 0-40 μM up to 100 times at a wavelength of 600nm. Measurements were carried out at a BD Fortessa flow cytometer. TN-XXL expression was checked assessing positive fluorescence in a 525±25 nm channel after 488nm excitation on a MACSQuant flow cytometer.

### Perfusion chamber

All *in vitro* experiments were carried out in Krebs-Ringer solution containing 6mM Ca^2+^ at 37°C. Cells were stimulated with anti-mouse IgM-F(ab)_2_ (Southern Biotech), ionomycin (4μM, Sigma), anti-CD40 antibody (xy), LPS (20 μg/ml, Sigma) or CpG (10 μg/ml, TIB Molbiol Berlin). Cell culture imaging experiments with ionomycin stimulation were performed using an open perfusion chamber system. Buffer solution was pumped through the heated chamber containing a poly-D-Lysin coated glass slide on which freshly and sterile isolated YellowCaB cells were grown for approx. 1 h. Ionomycin was added in the flow-through buffer supply. The lag time for the volume to arrive at the imaging volume was determined for each set-up and considered for analysis of ΔR/R over time. Anti-IgM-F(ab)_2_ antibody was given directly to cells within the open chamber in between acquisition time points. To visualize the reversibility of the sensor despite antibody still present, the experiment was performed in an open culture system without media exchange through a pump. To analyze if YellowCaB cells could repeatedly be stimulated, experiments were performed under continuous perfusion. Buffer flow was switched off with stimulation for several minutes and switched on again to dilute antibody out again for a second stimulation.

For analysis, regions of interest were determined based on randomly chosen single cells. Intensity density values of the raw citrine signal were divided by the intensity density values of the raw eCFP signal and related to the baseline ratio of the signals before stimulation.

### Cell transfers, immunization and surgical procedures

B cells from spleens of YellowCaB mice were negatively isolated using the Miltenyi murine B cell isolation kit via MACS. 5×10^6^ cells were transferred to a host mouse with a transgenic B cell receptor specific for an irrelevant antigen (myelin oligodendrocyte glycoprotein). B cells from spleens of B1-8^hi^:YellowCaB mice were transferred to WT C57/Bl6 mice. Host mice were immunized in the right footpad with 10 μg NP-CGG in complete Freund’s adjuvant 24h after B cell transfer. After six to eight days p.i., FDCs were labeled with Fab-Fragment of CD21/35-Atto590 or CD21/35-Alexa647 (inhouse coupling) into the right footpad. 24h after labelling, the popliteal lymph node was exposed for two-photon-imaging as described before^23^. Briefly, the anesthetized mouse is fixed on a microscope stage custom-made for imaging the popliteal lymph node. The mouse is shaved and incisions are made to introduce fixators that surround the spine and the femoral bone. The mouse is thus held in a planar position to the object table. The right foot is fixed by wire allowing to increase the tension on the leg to position the lymph node parallel to the imaging set up. A small incision is made to the popliteal area. The lymph node is exposed after freeing it from surrounding adipose tissue. A puddle around the lymph node is formed out of insulating silicon compound, then filled with NaCl and covered bubble-free with a cover slide.

### Intravital and live cell imaging and image analysis

Imaging experiments of freshly isolated B cells were carried out using a Zeiss LSM 710 confocal microscope. Images were acquired measuring 200-600 frames with 1 frame/3s frame rate while simultaneously detecting eCFP and citrine signals at an excitation wavelength of 405 nm.

For intravital two-photon ratiometric imaging, Z-stacks were acquired over a time period of 30-50 min with image acquisition every 30 seconds. eCFP and citrine were excited at 850nm by a fs-pulsed Ti:Sa laser and fluorescence was detected at 466±30 nm or 525±25 nm, respectively. Fluorescence signals of FDCs were detected in a 593±20 nm channel. For experiments including macrophage-staining, the fluorescence data has been unmixed for a possible overlap of the TN-XXL-citrine signal with that of the injected marker to prevent falsepositive colocalization analysis between the red efluor660 coupling of anti-CD169 and the green fluorescence of TN-XXL in the 525±25nm channel^68^.

For intravital FLIM experiments, eCFP fluorescence lifetime was measured with a time-correlated single-photon counting system (LaVision Biotec, Bielefeld, Germany). The fluorescence decay curve encompassed 10 ns with a time resolution of 55 ps and is a biexponential function containing the two mono-exponential decays of unquenched CFP and of FRET-quenched CFP, respectively.

### Analysis of two-photon data

For ratiometric analysis of two-photon data (here the exciting wavelength of 850 nm is also stimulating citrine fluorescence directly), fluorescence signals were corrected for spectral overlap (the eCFP to citrine ratio in 525±25 nm channel is 0.52/0.48) and refined by taking into account the sensitivity of photomultiplier tubes (PMTs, 0.37 for 466±30 nm and 0.4 for 525±25 nm). Ratiometric FRET for in vivo experiments was accordingly calculated as

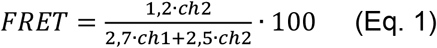

Evaluation of FLIM data was performed using the phasor approach^14,24^. Briefly, the fluorescence decay in each pixel of the image is Fourier-transformed at a frequency of 80 MHz and normalized, resulting into a phasor vector, with the origin at (0|0) in a cartesian system, pointing into a distinct direction within a half-circle centered at (0.5|0) and a radius of 0.5. For pure substances, lifetime vectors end directly on the half-circle, for mixtures of two on a connecting line between the respective pure lifetimes and within a triangle, if three substances are present, and so on. The distance between several fluorescence lifetimes on the half-circle is naturally distributed logarithmically, with the longest lifetimes closer to the origin. In the case of TN XXL, the extremes are the unquenched CFP fluorescence (2300 ps) and the CFP fluorescence completely quenched by FRET (700 ps). The location of the measured lifetime on the connecting line can directly be translated into the amount of either CFP state and thus to a corresponding calcium concentration.

Due to the scattering environment in tissue, the FLIM-signal of the donor is shifted towards the (0/0) position in the phasor plot, indicative for the infinite lifetime of noise. Thus the phase vectors are shortened and end within a triangle between the points (0|0), approximately 2300 ps and 700 ps. The noise should therefore be considered as a contribution of a third exponential component. In a non-fluorescent medium, we measured a noisy FLIM-signal under similar experimental conditions as those used in the intravital experiments and Gaussian-fitted the histograms of real and imaginary part after the transformation to the frequency domain. The Gaussian fit of each part gives a mean, which indicates the center of the noise distribution, as wells as the width 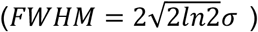, which was the same for both parts (real and imaginary axis) and gives us the radius (½FWHM = 0.2 (green, solid arc in Fig. S5) within which we expect only noise, i.e. the signal-to-background-ratio is unreliable small. In order to increase the accuracy, we enlarged the radius to ¾FWHM = 0.3 (green, dashed arc in Fig. S5) and all data points within that radius were excluded from further analysis. A second filtering was applied by determining the signal-to-noise ratio (SNR) from the summed TCSPC signal for all segmented cells. In Imaris, a sphere of radius=20μm was determined around each cell to establish a reference value for background noise. The SNR was then calculated as follows (pixel by pixel):

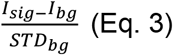

With I_sig_ being the intensity of TCSPC signal, I_bg_ the intensity of background noise and STD_bg_ the background noise standard deviation^69^ AG specific signals with SNR<2 and non-Ag specific signals with SNR<1 were excluded from the analysis. All other phasor data points were projected on the segment connecting FRET-quenched and unquenched CFP fluorescence lifetime, in order to determine absolute calcium concentrations as follows:

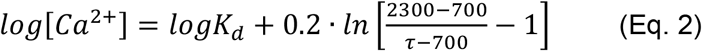

with Kd = 453 nM calcium^15^. The linear range of the TN-XXL titration curve was determined to be 265nM-857nM. All values below or above these margins are subject to uncertainty and therefore simply referred to as <265nM or >875nM, respectively.

### Statistical information

Time dependent FRET curve analysis shows representative graphs for the number of analyzed cells and independent experiments given. For multiple curve analysis, mean is shown and SD indicated in each data point. For column analysis, One-way ANOVA with Bonferroni Multiple Comparison Test was applied with a confidence Interval of 95%.

### Data availability

All raw data and analyzed data shown here are stored on institutional servers and may be accessed upon request to the corresponding author.

### Code availability

Python-based code for phasor analysis can be provided upon request.

## Acknowledgements

We thank Patrick Thiemann, Vivien Theissig and Manuela Ohde for animal caretaking. We thank Robert Günther for excellent surgical assistance and Peggy Mex for cell isolations and stainings. We thank Ralf Uecker for microscope facility services. Further thank goes to Mathis Richter for his help with SNR exclusion analysis and proofreading of the manuscript. This study has been supported by the DFG grant TRR130.

## Author contributions

A.E.H, R.A.N. and C.U. designed the study and single experiments. F.K. kindly provided mice. C.U. and A.R. conducted experiments. R.L., A.R. and R.A.N. developed mathematical analysis strategies and provided bioinformatical support. L.N. supported with expertise in flow cytometric calcium flux measurements. H.R. provided help for general approach of GECI experiments. C.U., R.L., A.E.H. and R.A.N wrote the manuscript.

## Additional information

Supplementary information is available for this paper.

Competing interest: The authors declare no competing financial interests.

Correspondence: All requests for additional information and material should be addressed to.

Reprints and permission information is available online at:

A preprint of this paper is available at bioRxiv: https://doi.org/10.1101/2019.12.13.872820

