## Supplementary figures S1-S6 for "Intravital quantification of absolute cytoplasmic B cell calcium reveals dynamic signaling across B cell differentiation stages"

**Supplementary figure S1**

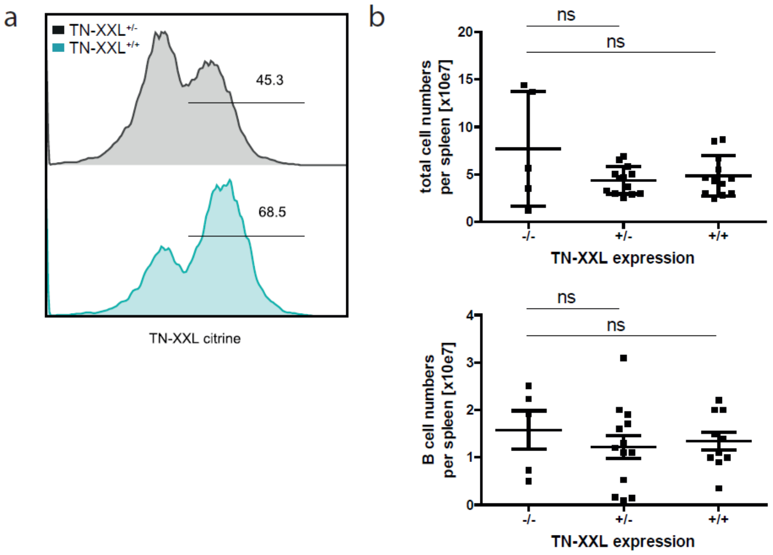

#### Supplementary figure S2

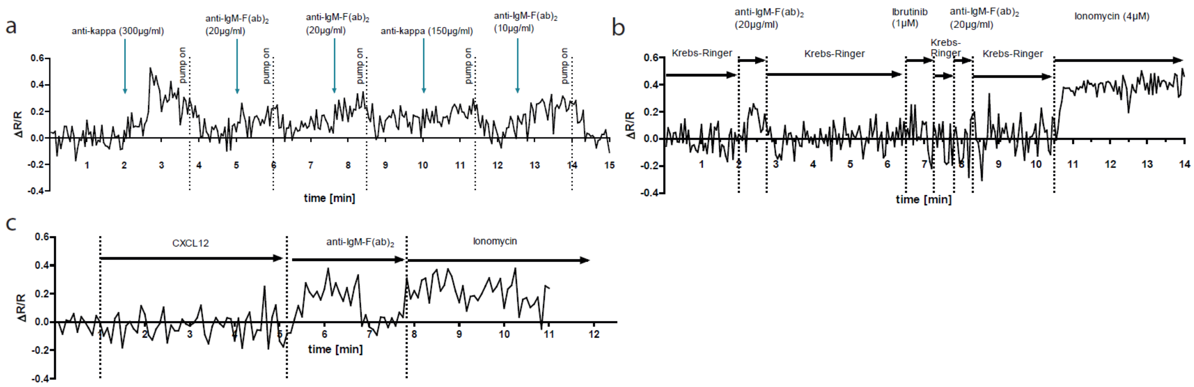

**Supplementary figure S3**

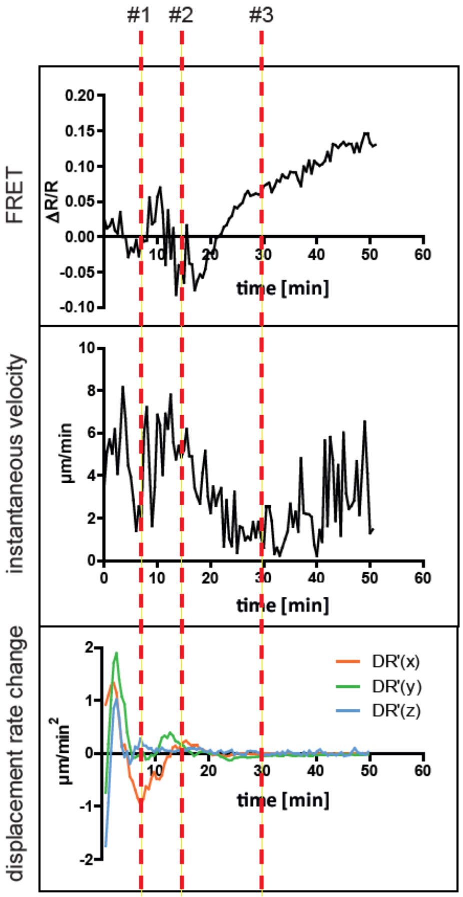

**Supplementary figure S4**

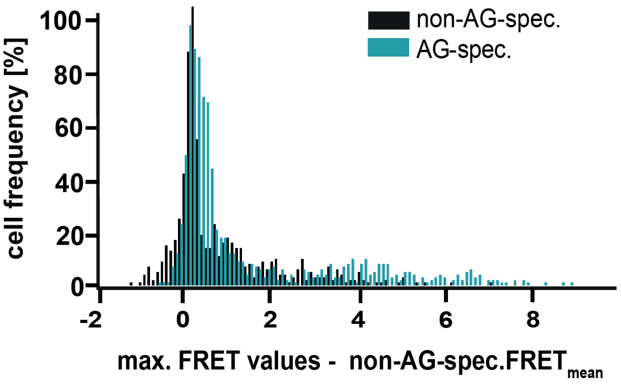

### Supplementary figure S5

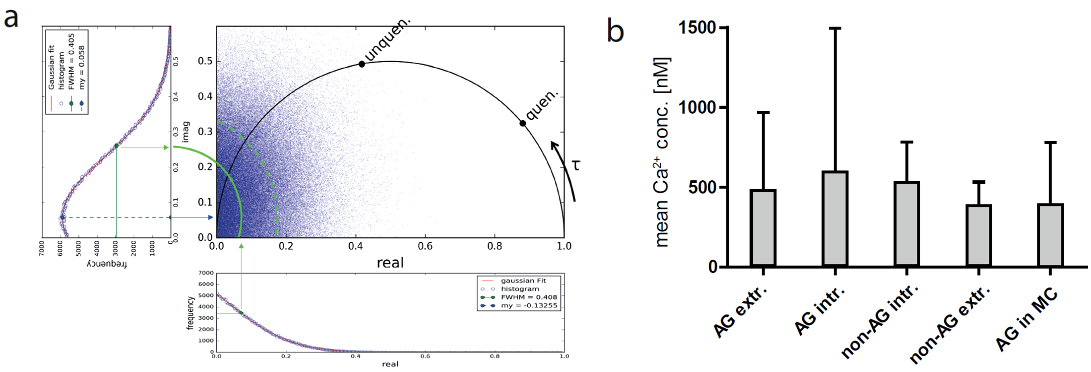

**Supplementary figure S6**

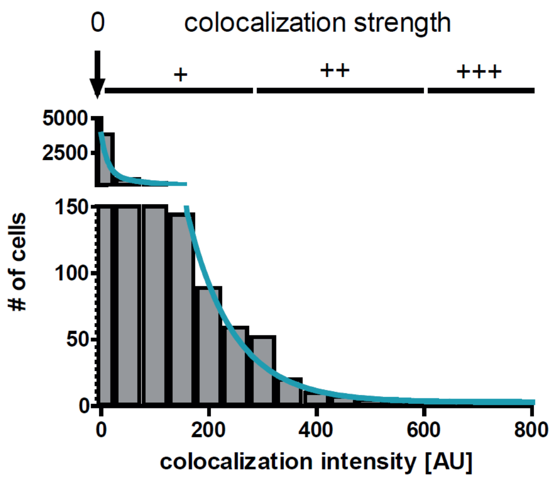
