## Supplementary figure legends for "Intravital quantification of absolute cytoplasmic B cell calcium reveals dynamic signaling across B cell differentiation stages"

Supplementary figure captions:

**Fig S1:** Genotyping of YellowCaB mice and cell numbers. a Exemplary flowcytometric analysis of YellowCaB splenocytes pregated for CD19^+^ in 525±25nm channel after 488nm excitation. Half of CD19^+^ B cells from heterozygous (TN-XXL^+/-^) mice and two thirds of CD19^+^ B cells from homozygous (TN-XXL^+/+^) mice show positive signals in the TN-XXL gate (horizontal line). b Total cell numbers (top) and B cell numbers (bottom) of YellowCaB and wildtype mice. Different ages and sex pooled.

**Fig S2:** Cytosolic calcium concentrations change detectable with TN-XXL can be provoked up to five times and is BCR specific. Confocal measurement and plot of ΔR/R over time. a In an open perfused system (Krebs-Ringer solution 6mM Ca^2+^), single cells were stimulated with indicated concentrations of BCR anti-light- or anti-heavy chain antibody (arrows). Antibody was added directly to the imaging chamber volume. Perfusion pump was switched off with stimulation, and switched on again at indicated time points (dashed lines). b Open perfusion of single cells with 6mM Ca^2+^ Krebs-Ringer solution or 6mM Ca^2+^ Krebs-Ringer plus reagents indicated. Here, stimulation was performed under continuous flow via changing the reservoir of the afferent buffer solution. c Stimulation of single cells with cytokine CXCL12. No ΔR/R signal change is detectable after cytokine exposure, but after BCR stimulation. Experimental set-up as described in b.

**Fig S3:** Cell-to-cell contacts coincide with FRET signal change and altered cell motility. Intravital imaging parameters of GC YellowCaB cell making multiple contacts with FDCs (#1-3, red dashed lines). Top: ΔR/R FR ET signal change of tracked cell over time, first 5 time point were taken as baseline. Middle: Instantaneous velocity over time. Bottom: First derivative of displacement rate in direction of X (orange), Y (green) and Z (blue). All DR’=0 means complete halt of cell dislocation.

**Fig S4:** Relative comparison of AG- and non-AG-specific YellowCaB cells in ratiometric intravital imaging in lymph nodes. Mean non-AG-spec. FRET was chosen to normalize values across five different measurements. AG-spec cells (blue) show extra population not visible among non-AG-spec. cells at value 3.5-8 AU.

**Fig S5:** Absolute quantification of cytosolic calcium concentration in YellowCaB cells. a Determination of noise exclusion radius for phasor analysis. b mean calcium concentration and variances (SD) among B cell populations *in vivo*.

**Fig S6:** Colocalization histogram and exponential fit for analysis of colocalization between CD169^+^ macrophages and extrafollicular YellowCaB cells. Colocalization was intensity was determined via setting Intensity thresholds for red (macrophages) and green (YellowCaB cells) and calculation of overlap in each pixel. Colocalizaiton of <1 was considered non-colocalized, colocalization intensity >1 was considered colocalized with weak (+), Intermediate (++) or strong (+++) contact depending on decay rates (<25% for ++, <10% for +++ contact) of biexponential fit.

**Supplementary movies**

**Movie S1:** Detail of intravital ratiometric imaging (d 8p.i.) within GC. Non-AG-specific YellowCaB cells had been transferred to a restricted host, FDCs were *in vivo*-labeled with anti-CD21/35 antibody (white). 3D-surface rendering and single-cell tracking (track line in yellow) with color coding ranging from blue= low ΔR/R to red=high ΔR/R. 103 frames, 7fps, scale bar 50µm.

**Movie S2:** Side-by-side depiction of fluorescence, FLIM and cell-based phasor data of intravitally imaged GC (d 8p.i.), single z-plane. Left: Fluorescence data with AG-specific YellowCaB cells (green) and stained non-AG-specific YellowCaB cells transferred one day prior to imaging (red; autofluorescence of capsule also visible in the same channel). 4 frames per second (fps), scale bar 50µm. Middle: false color-coded presentation of fluorescence lifetime tau (0-3000ps, see range scale in Fig 4b). 4 fps, scale bar 50µm. Right: Raw cell-based phasor plot with cells segmented according to fluorescence and spatial distribution, subsets indicated. 4fps.

**Movie S3:** Detail of movie S2 within MC**,** side-by-side depiction of fluorescence and FLIM data, single z-plane. Left: Fluorescence data with AG-specific YellowCaB cells (green) and autofluorescence of capsule (red). 4 fps, scale bar 50µm. Right: false color-coded presentation of fluorescence lifetime tau (0-3000ps, see range scale in Fig 4b). 4 fps.

**Movie S4: 3D projection** Intravital imaging of GC and subcapsular sinus. YellowCaB cells (green, with track lines) and SCSM (red), stained by CD169 *in vivo*-labeling. White circles highlighting AG-specific B cells migrating along subcapsular space. 4fps, scale bar 50µm.
